# Sulfotransferase 4A1 (SULT4a1): A Novel Neuroprotective Protein in Stroke

**DOI:** 10.1101/2025.04.07.647574

**Authors:** M Iqbal Hossain, Junaid Khan, Jun Hee Lee, Charles N Falany, Peter H King, Mark Boulding, Shaida A Andrabi

## Abstract

SULT4a1, a member of the cytosolic sulfotransferase family, is predominantly expressed in neurons and plays potentially vital roles in regulating neural survival and function. SULT4a1 protects against mitochondrial dysfunction and oxidative stress. SULT4a1 levels decrease in experimental stroke models and may play a critical neuroprotective role in mitigating neuronal injury caused by oxygen-glucose deprivation (OGD) and ischemic stroke, as shown in a transient middle cerebral artery occlusion (tMCAO) mouse model. In this study, we investigated the neuroprotective role of SULT4a1 in OGD and tMCAO and highlighted its expression pattern and involvement in maintaining mitochondrial function and reducing oxidative stress, two early pathophysiological features in stroke and related neuronal injury. Our data show that decreased SULT4a1 expression in OGD conditions and in the tMCAO mouse brain leads to enhanced neuronal damage, emphasizing the importance of SULT4a1 in preserving neuronal integrity. Loss of SULT4a1 alone was sufficient to decrease mitochondrial function in mouse cortical neurons. Notably, overexpression of SULT4a1 preserved mitochondrial function, reduced the loss of mitochondrial membrane potential, and diminished oxidative stress, as evidenced by lower reactive oxygen species (ROS) production and reduced protein carbonylation. These results indicate that modulating SULT4a1 expression in stroke could offer a promising strategy for preventing neuronal damage. Indeed, overexpression of SULT4a1 via stereotaxic injection of AAV9 into the mouse brain mitigated tMCAO-related brain injury and functional deficits over time. The findings of this study indicate that SULT4a1 may protect neurons in stroke and related brain injury, possibly by maintaining mitochondrial function and redox homeostasis through mechanisms that are still unknown. It is likely that SULT4a1 regulates neuroprotective processes in both the mitochondria and the cytosol. However, further research is needed to clarify the specific molecular pathways involved in its neuroprotective function.

## Introduction

Stroke is a major cause of death and disability, characterized by complex pathophysiology and limited therapeutic options.^1,2^ The disruption of blood flow leads to a deficiency in oxygen and glucose, which results in mitochondrial dysfunction, bioenergetic failure, and the subsequent dysregulation of cellular calcium and redox balance.^3,4^ These early pathological events contribute significantly to neuronal loss and brain damage following a stroke^4^. Strategies aimed at preserving mitochondrial function and redox homeostasis have demonstrated neuroprotective effects in experimental stroke models.^5,6^ Our previous work has shown that Sulfotransferase 4A1 (SULT4A1), a member of the cytosolic Sulfotransferases (SULT) enzyme family, is localized to mitochondria and regulates mitochondrial function and redox balance in both neuronal and yeast models of oxidative stress.^7,8^ These studies indicate that SULT4A1 may offer neuroprotection in stroke through its influence on mitochondrial function and cellular homeostasis. While SULT4A1 shares limited sequence similarity with other SULT enzymes, it is highly conserved across species, with over 98% sequence identity, suggesting essential functions in a broad range of organisms.^9^ SULTs are Phase II metabolic enzymes that transfer a sulfonate (-SO3-) group from 3′-phosphoadenosine 5′-phosphosulfate (PAPS) to various substrates, converting hydrophobic compounds into more water-soluble metabolites, thereby modulating their pharmacological and toxicological activities.^10,11^ However, no sulfonation activity or specific substrates for SULT4A1 have been identified to date.^12^ Structural analysis of SULT4A1 reveals a shortened enzyme-binding loop and a lack of the conserved lysine residue essential for PAPS binding, which suggests that SULT4A1 may not function as a classical sulfonating enzyme.^12^ While it is assume that SULT4A1 might be catalytically inactive, further studies are necessary to confirm this assumption. It is also possible that SULT4A1 regulates the activity of specific proteins through direct physical interactions, independent of its enzymatic function.

SULT4A1 is uniquely expressed in neurons of the brain, contrasting with other SULT family members.^13,14^ This high conservation across species and its neuron-specific expression suggests that SULT4A1 may have distinct biological roles in the brain. Sult4a1 knockout mice display a significant neurological impairments, including tremors, anxiety, and reduced body size, highlighting the importance of SULT4A1 in the brain.^15^ Recent research has underscored role of SULT4a1 in synaptic development, synaptic maturation, and synaptic plasticity.^16^ Reduced expression of SULT4A1 has been observed in the cortices of individuals with autism spectrum disorder,^17^ and SULT4A1 haploinsufficiency has been implicated in Phelan-McDermid Syndrome (PMS) and depression.^18^ Additionally, SULT4A1 variants are associated with the genetic underpinnings and psychopathology of schizophrenia, as well as response to antipsychotic treatments.^19^ Collectively, these findings emphasize a critical role of SULT4A1 in maintaining neuronal health, with alterations in its expression leading to disrupted brain processes.

A recent study showed a reduction in SULT4A1 levels in mice subjected to middle cerebral artery occlusion (MCAO)^20^. Our data also show that SULT4a1 levels decrease in oxygen glucose deprivation (OGD) in mouse neuronal cultures and in the brain of MCAO mice. Given its pivotal role in regulating mitochondrial function and oxidative homeostasis during oxidative stress^7^, it remains unclear whether the loss of SULT4A1 contributes to mitochondrial dysfunction and redox imbalance in stroke. Furthermore, it is unknown whether restoring SULT4A1 levels could offer neuroprotection in models of stroke. To address this, we employed viral-mediated overexpression of SULT4A1 and demonstrated that SULT4A1 confers neuroprotection in stroke, suggesting a therapeutic potential of SULT4a1 in stroke and related brain injury.

## Result

### 1. SULT4a1 is a neuron-specific protein

SULT4a1 is a brain-specific protein, and the knockout mice display severe phenotype including smaller size and tremor^7,15^, suggesting vital roles of SULT4a1in the brain. Brain-specific expression of SULT4A1 in has been documented in previous studies^15^; however, a comprehensive understanding of its expression within the brain has been limited. We performed immunostaining to visualize the presence of SULT4A1 protein across various brain regions. Our results revealed a strong and consistent signal in all major brain regions, including the cortex, hippocampus, cerebellum, and basal ganglia (Fig 1A). We also utilized western blotting as a complementary assay to assess SULT4A1 expression in tissue samples from different brain regions. These western blot results corroborated the immunohistochemical findings, demonstrating high levels of SULT4A1 across a range of brain regions (Fig 1B). Together, these methods strengthen the evidence for the widespread expression of SULT4A1 in the brain, suggesting its potential importance for proper brain health and function.

**Figure 1.**
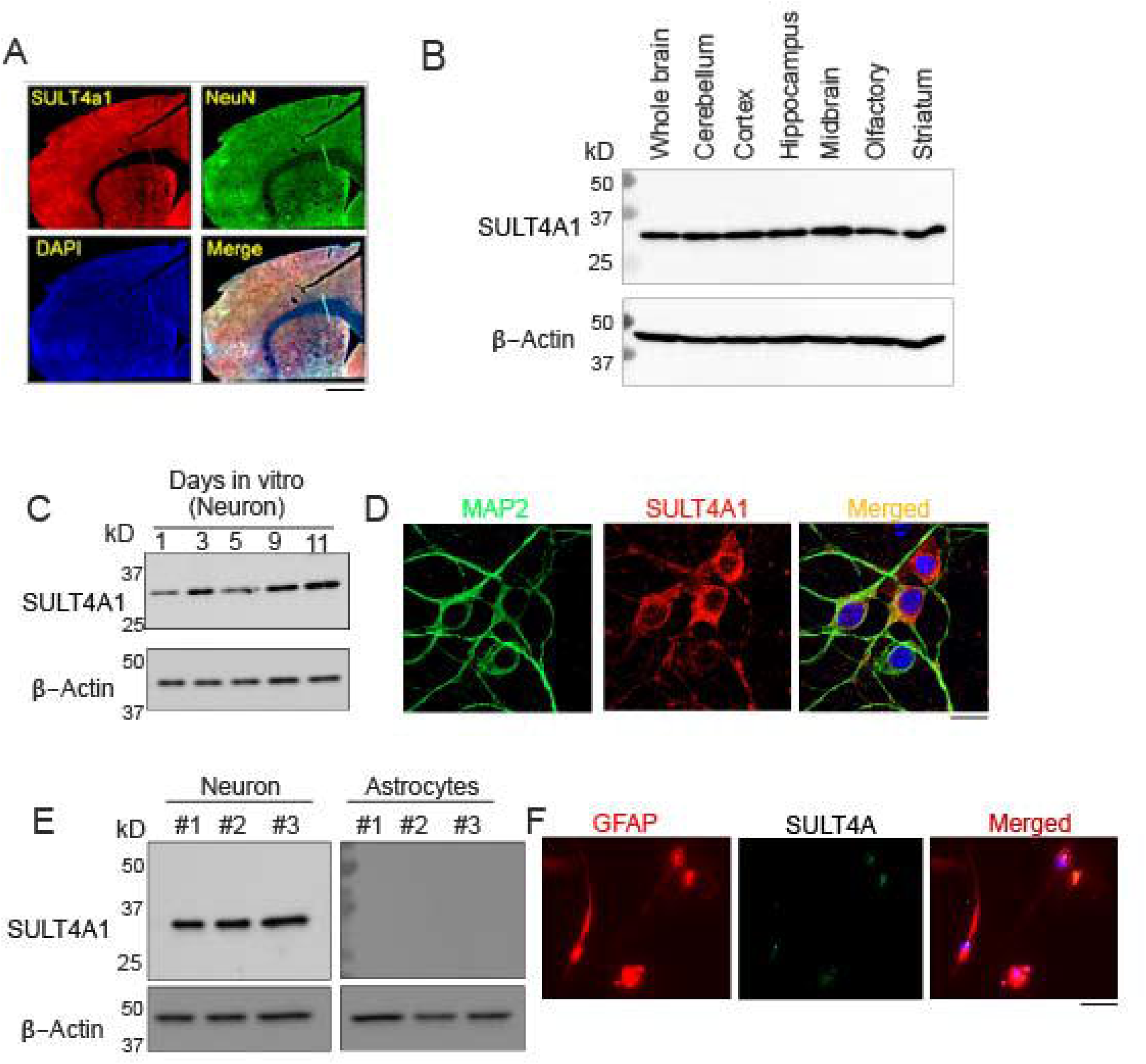
Expression of SULT4A1 in the brain and primary neurons. **A.** Confocal image of a brain section showing expression of SULT4A1 in cortex and striatum. NeuN is used as a marker of neurons. Scale bar: 20 μm. **B.** Representative western blot showing expression of SULT4A1 in different parts of a mouse brain. β-Actin is used as a loading control. **C.** Representative western blot showing expression levels of SULT4A1 in primary neurons at different days of (DIV1 to DIV 11) culture. **D.** Confocal images of primary neurons showing neuron specific expression of SULT4A1. MAP2 is used as a neuronal marker. Scale bar: 20 μm. **E.** and Western blot **(E)** and Confocal images **(F)** showing exclusive expression of SULT4A1 in neurons, not in Astrocytes. GFAP is used as the astrocytic marker. Scale bar: 20 μm.

Further analysis of SULT4A1’s cellular localization using markers for astrocytes, microglia, and neurons revealed that SULT4A1 is exclusively expressed in neurons (Fig 1C & D), with no detectable levels in astrocytes or microglia (Fig 1E & F). These findings imply that SULT4A1 is an essential neuronal protein potentially required for the regulation of key neuronal functions and survival pathways, through mechanisms that are yet to be fully understood.

### 2. Protein levels of SULT4a1 decrease in stroke

SULT4a1 is a neuron-specific protein with some vital but yet unknown brain functions^7^. A recent study revealed that SULT4a1 expression decreases in a mouse stroke model. ^20^ We assessed the level of SULT4a1 in neurons following OGD using Western blot and immunocytochemistry. We used a 60-min OGD followed by oxygen and glucose resupply (OGR) in mouse cortical neurons, as outlined in Figure 2 (Fig 2 A & B). Western blotting results showed that the Sult4a1 protein levels decrease in OGD-exposed neurons at 1 h, 3 h, and 6 h after the termination of OGD (Fig 2 C & D) compared to control neurons. These data indicate that loss of SULT4a1 may play a pathophysiological role in stroke.

**Figure 2.**
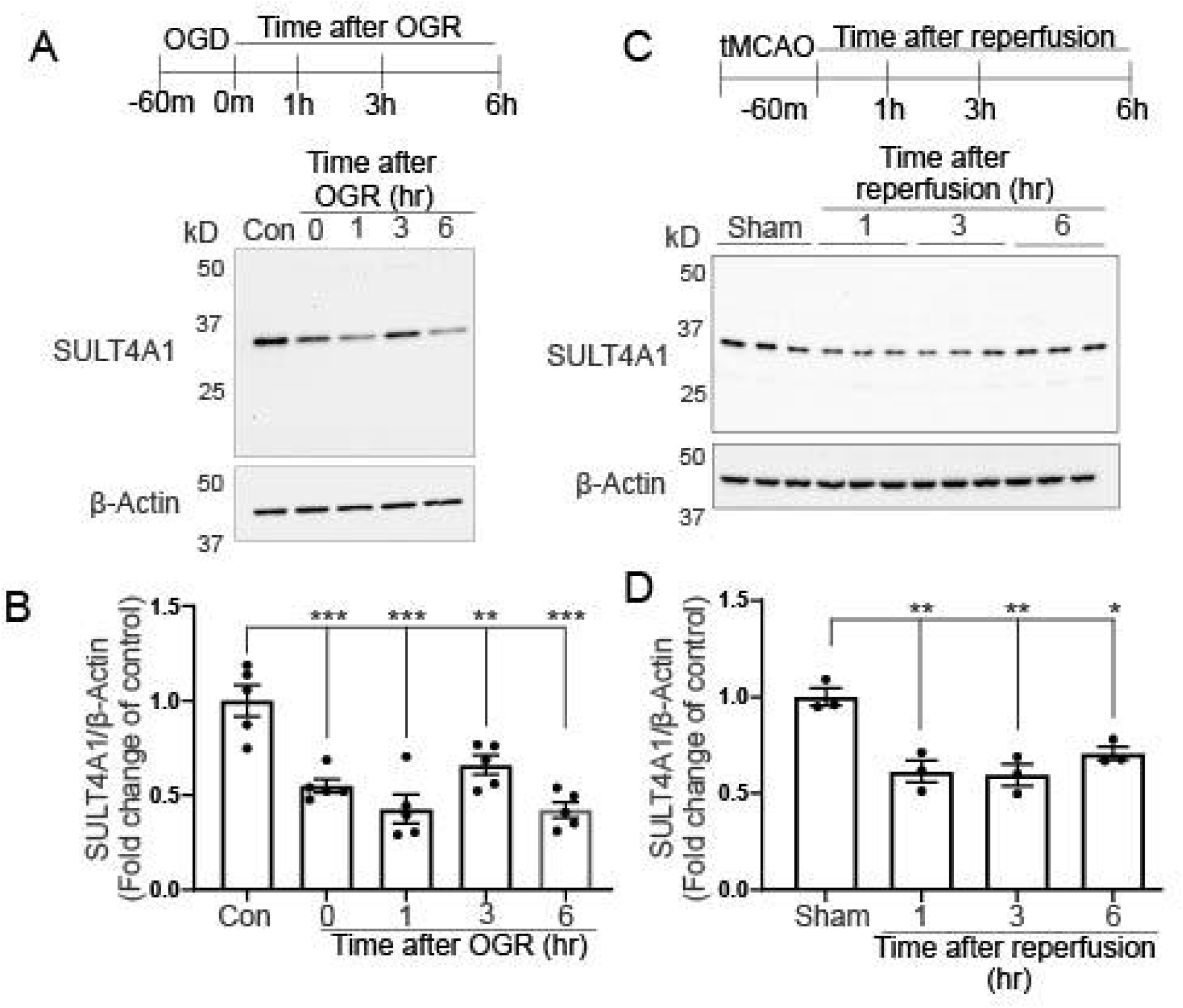
Level of SULT4A1 OGD neurons and stroke brain. Top: Schematic diagram showing the experimental design of OGD/R to neurons. Bottom: Representative western blot **(A)** and Quantification **(B)** showing expression levels of SULT4A1 in OGD neurons at different time points of OGR (n = 5). **C.** Top: Schematic diagram showing MCAO and reperfusion time in mouse brain. 1, 3, and 6 hr indicate brain collection periods. Bottom: Representative western blot showing expression of SULT4A1 in sham and MCAO brain at different time points of reperfusion. **D.** Quantification of expression levels of SULT4A1 obtained from (C). β-Actin is used as a loading control. Data are mean ± SEM (n = 3). *P < 0.05, **P < 0.01, ***P < 0.001 vs indicated groups, calculated with two-way ANOVA followed by Tukey’s post hoc test.

### 3. Loss of SULT4a1 results in mitochondrial dysfunction in mouse cortical neurons

SULT4a1 is primarily expressed in neurons, and it is not present in cancerous or dividing cells. Our previous work demonstrated that ectopic expression of SULT4a1 in SH-SY5Y cells protects them from oxidative stress-induced mitochondrial dysfunction, with some SULT4a1 localized to the mitochondria.^7^ To determine whether the loss of SULT4a1 alone is sufficient to cause mitochondrial defects in mouse primary cortical neurons, we used shRNA to knock down SULT4a1 via lentiviral transduction at Day in Vitro (DIV) 5. By DIV 11, we confirmed efficient knockdown of SULT4a1 (Fig 3A). To evaluate mitochondrial function, we measured the Oxygen Consumption Rate (OCR) via an XFe Seahorse Flux analyzer on DIV 11. The neuronal media was replaced with Seahorse media containing 10 mM glucose, 1 mM glutamine, and 1 mM pyruvate. Basal respiration, ATP turnover, and maximal respiration were assessed by sequentially adding oligomycin (1 µM), CCCP (1 µM), and rotenone/antimycin A (1 µM each). Our results showed that SULT4a1 knockdown significantly impaired mitochondrial function in mouse cortical neurons (Fig 3B) Specifically, basal respiration (Fig 3C), maximal respiration (Fig 3D), and ATP turnover (Fig 3E) were all reduced in neurons with SULT4a1 knockdown. These findings suggest that the loss of SULT4a1 may contribute to mitochondrial dysfunction in stroke, raising the question of whether restoring SULT4a1 through overexpression could protect against stroke-induced neuronal loss and mitochondrial dysfunction.

**Figure 3.**
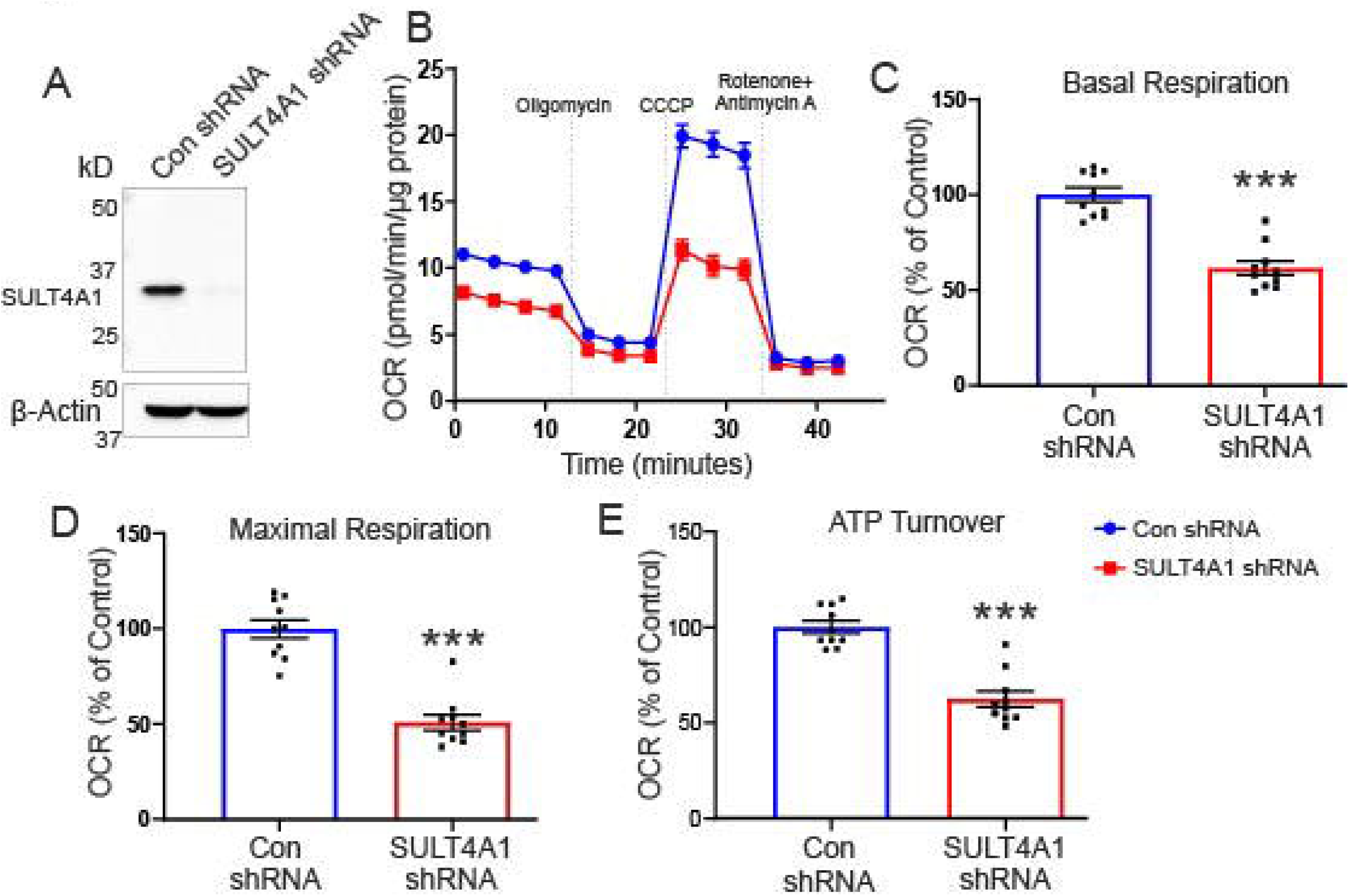
Assessment of mitochondrial function in SULT4A1 KD neurons. **A.** Western blot showing depletion (>95%) of SULT4A1 levels in SULT4A1 shRNA lentivirus transduced neurons. **B**. Representative OCR in neurons transduced with control shRNA or SULT4A1 shRNA lentivirus. Lentivirus was added to neurons on DIV 5, and OCR was measured at DIV 11. **C.** Basal respiration, **D.** Maximal respiration, and **E.** ATP turnover. All these parameters were calculated relative to the Control shRNA transduced neuron. Data are mean ± SEM (n = 10). ***P < 0.001 vs indicated groups, calculated with two-way ANOVA followed by Tukey’s post hoc test. OCR experiments were repeated three times with similar results.

### 4. SULT4a1 improves mitochondrial function in neurons exposed to OGD

Previously, we demonstrated that SULT4a1 protects against oxidative stress-induced mitochondrial dysfunction and redox imbalance in human neuronal cells^7^. Both mitochondrial dysfunction and disruption in oxidative homeostasis are early pathophysiological processes in stroke.^21,22^ We employed lentiviral-mediated overexpression of SULT4a1 or GFP control in mouse cortical neurons on DIV 5 (Fig 4A) and exposed these neurons to OGD on DIV 11 followed by analysis of mitochondrial function using a Seahorse Flux analyzer (Fig 4B). We assessed OCR as mitochondrial function at 3hr after the termination of OGD, and measured basal respiration, ATP turnover, and maximal respiration by sequentially adding oligomycin (1 µM), CCCP (1µM), and rotenone/antimycin A (1µM each). Our data revealed that SULT4a1 overexpression significantly increases the OGD-mediated loss of mitochondrial function as compared to OGD-exposed neurons transduced with GFP control lentivirus (Fig 4B). Mitochondrial parameters including basal OCR (Fig 4C), maximal OCR (Fig 4D) and ATP turn over (Fig 4E) were significantly preserved in OGD-exposed neurons with SULT4a1 overexpression. One of the indications of mitochondrial dysfunction is the loss of mitochondrial membrane potential (Δψm). We assessed Δψm in neurons at 1, 3, and 6 hr after using Tetramethylrhodamine ethyl ester (TMRE) live-cell imaging via an LSM 710 Confocal microscope (Carl Zeiss). Our data revealed that OGD results in a significant loss of Δψm assessed at 1, 3, and 6 hr after the termination of OGD (Fig 4 F,G,H). SULT 4a1 overexpression significantly preserved the Δψm in OGD-exposed neurons at all the observed time points (Fig 4 F,G,H). The data demonstrate that SULT4a1 preserves mitochondrial function in OGD-exposed neurons and maybe a drug target for recovering/improving mitochondrial function in stroke.

**Figure 4.**
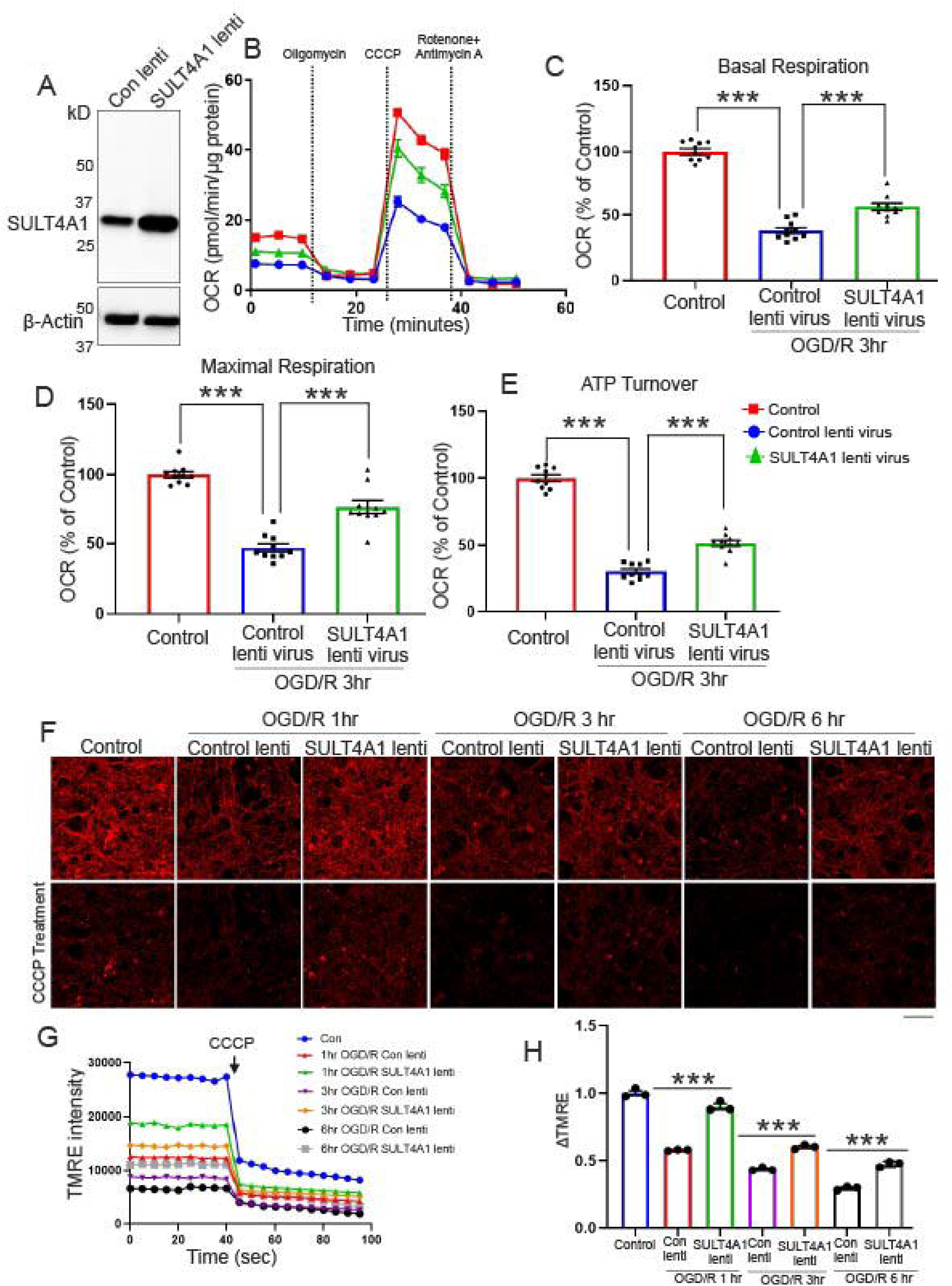
Assessment of mitochondrial function in OGD neurons transduced with SULT4A1 lentivirus. **A.** Western blot showing expression levels of SULT4A1 in neurons transduced with SULT4A1 virus. Data showing a ∼4-fold increase in the expression level of SULT4A1 **B**. Representative OCR in neurons transduced with control or SULT4A1 virus following exposure to OGD. OCR was measured at 3hr of OGR. Lentivirus was added to neurons on DIV 6, and OCR was measured at DIV 10. **C.** Basal respiration, **D.** Maximal respiration, and **E.** ATP turnover. All these parameters were calculated relative to the Control neuron. OCR experiments were repeated three times with similar results. Data are mean ± SEM (n = 10). **F**. Representative confocal images showing TMRE intensity in neurons transduced with control or SULT4A1 lentivirus following OGD. TMRE intensity was measured at 1, 3, and 6 hr of OGR. Scale bar = 20 μm. **G,** Time-lapse quantification of TMRE fluorescence intensity before and after CCCP addition. **H.** Bar graph showing quantification of TMRE intensity change (ΔTMRE) before and after CCCP addition in neurons shown in figure **(F).** Data are mean ± SEM (n = 3). ***P < 0.001 vs indicated groups, calculated with two-way ANOVA followed by Tukey’s post hoc test.

### 5. SULT4a1 overexpression protects against redox imbalance in OGD-exposed neurons

Mitochondrial dysfunction results in oxidative imbalance via excessive generation of reactive oxygen species (ROS).^23^ Protein carbonylation is a marker for oxidative stress and ROS production. We measured protein carbonylation in neurons after OGD via western blotting using an antibody specific to protein carbonyl. Our data show that protein carbonyl increases in control lentiviral expressing neurons following OGD, which is significantly reduced by SULT4A1 overexpression in mouse cortical neurons (Fig 5 A & B). We also assessed ROS generation using CellROX deep red live-cell imaging in control lentiviral transduced neurons and SULT4a1 overexpressing neurons at 1h, 3h and 6h following the termination of OGD. Our data indicate that OGD in control lentivirus expressing neurons causes an increase in ROS levels at all the observed time points after OGD termination (Fig 5 C & D) while SULT4a1 overexpression significantly decreases the ROS levels at all time points following OGD (Fig 5 C & D). These data demonstrate that SULT4a1 is a neuroprotective protein, and it acts by improving redox homeostasis in neurons.

**Figure 5.**
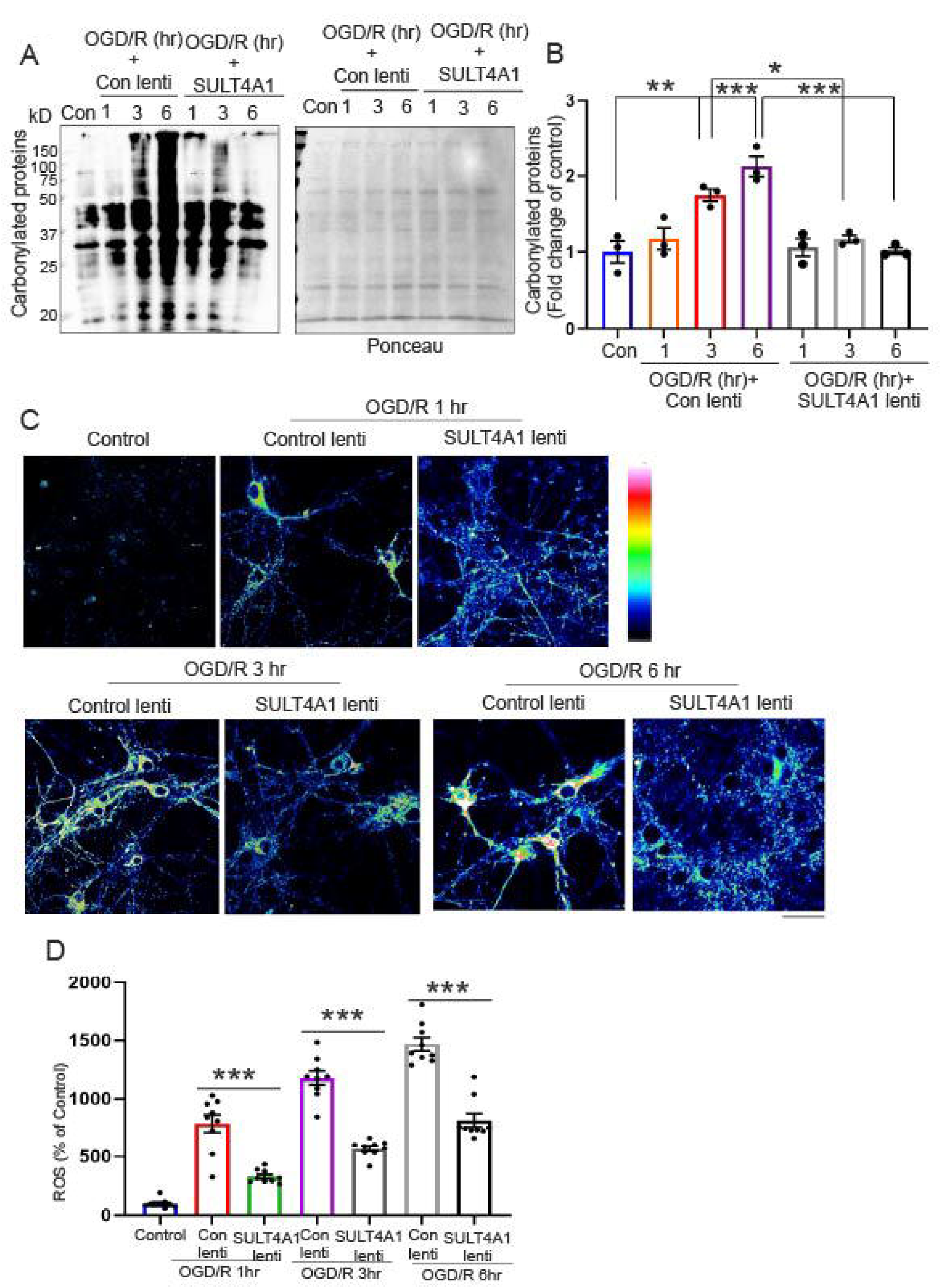
Assessment of the effect of SULT4A1 on the levels of reactive oxygen species. **A.** Representative western blot of protein carbonylation levels in control and OGD neurons transduced with control or SULT4A1 virus. Carbonylation levels were assessed at 1, 3 and 6 hr of OGR. Ponceau was used to show equal loading of proteins**. B.** Quantification of protein carbonylation levels in control and OGD neurons transduced with control or SULT4A1 lentivirus. Data are expressed as mean ± SEM. n=3 **C.** Representative confocal images of CellRox fluorescence intensity in control and OGD neurons transduced with either control virus or SULT4A1 virus. Measurements of CellRox intensity were performed at different time points of OGR as shown in the figure. Scale bar = 20 μm. **D.** Quantification of CellRox fluorescence intensity (ROS). Data are mean ± SEM. *P < 0.05, **P < 0.01, and ***P < 0.001 vs indicated groups, calculated with two-way ANOVA followed by Tukey’s post hoc test.

### 6. SULT4a1 is neuroprotective in OGD and reduces the brain infract volume in MCAO mice

We assessed whether SULT4A1 overexpression in mouse cortical neurons protects against OGD-mediated cell death. We transduced mouse primary cortical neurons with either control lentivirus or SULT4A1 lentivirus on DIV 5. We then exposed these neurons to 60 minutes of OGD on DIV 11 and assessed cell death using Alamar Blue 24 hours after the termination of OGD. Our data show that SULT4A1 overexpression protects against OGD-mediated cell death in mouse cortical neurons (Fig 6A). To assess the neuroprotective role of SULT4a1 *in vivo*, we overexpressed SULT4a1 in the mouse brain via stereotaxic injection of AAV9 driven by Syn1 promotor. We use GFP AAV9 as control as control. We injected AA9 (1µl × 10^13^) into two different locations in the MCA territory to ensure entire brain region around MCA that is affected by MCAO is transduced by the virus. Three weeks after AAV injections, we assessed SULT4a1 overexpression in a cohort of mice (n=3) via western blots. Our data show that there was a drastic increase in SULT4a1 in mouse brain following AAV9 injections (Fig 6 B & C). After three weeks of AAV infection, the mice were subjected to stroke surgeries via tMCAO using 45 min occlusion followed by reperfusion. In a subgroup of animals TTC staining was used to measure the infarct volume 24h after the reperfusion. Our data show that SULT4a1 significantly reduces the brain infarct volume in MCAO mice (Fig 6 D & E). Infract volume analysis using longitudinal MRI at day 3, day 15 and day 30 after reperfusion revealed that infarct volume gradually decreases spontaneously with time in control GFP expressing mice (Fig 6F). However, SULT4a1 expressing mice show significantly smaller infarcts at all the assessed time points in comparison to control GFP expressing MCAO mice (Fig 6F)

**Figure 6.**
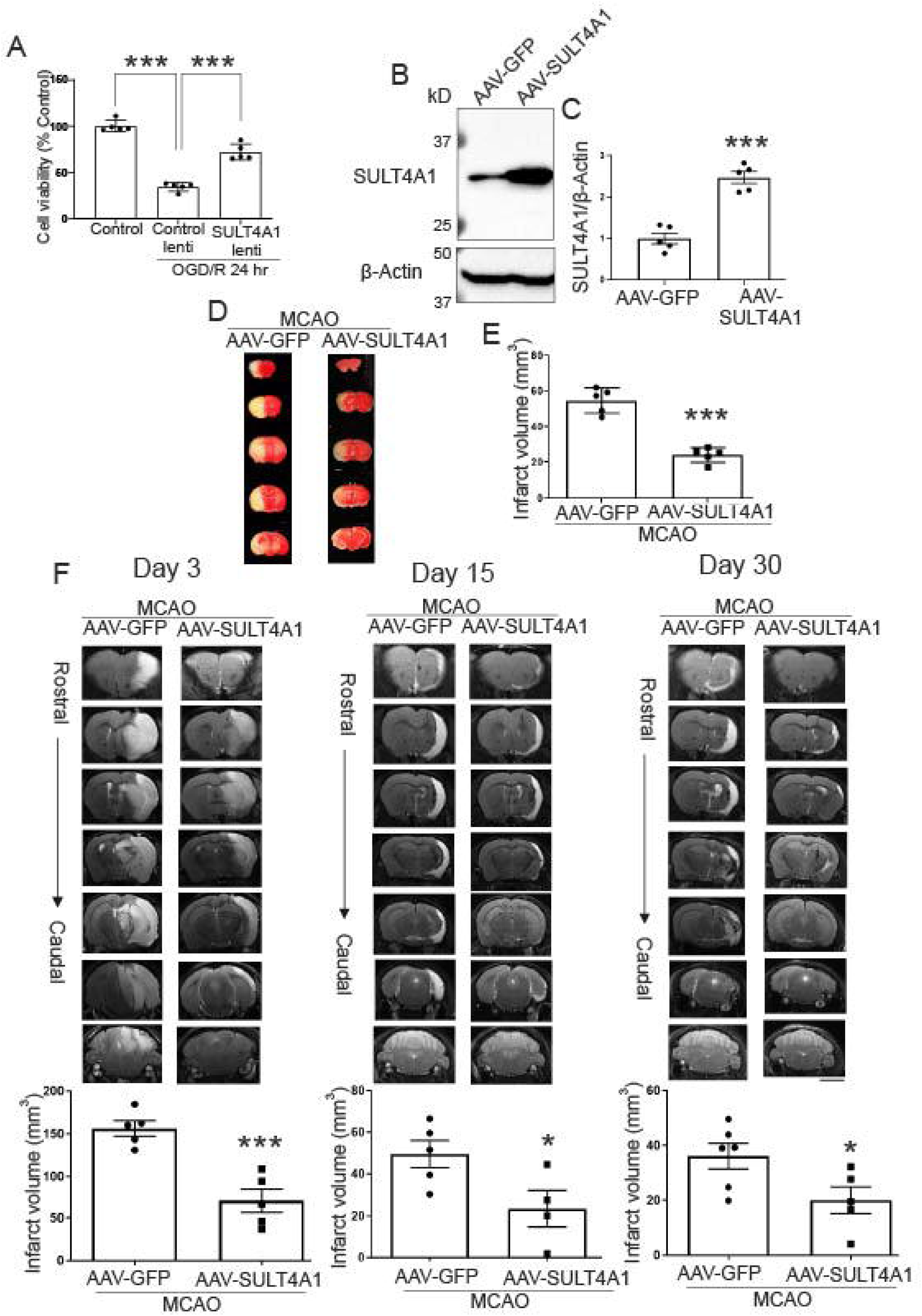
Assessment of the protective effect of SULT4A1 **A.** Cell viability (Alamar blue) in control and OGD neurons transduced with control or SULT4A1 lentivirus. Neurons were transduced with viruses at DIV 5 and exposed to OGD at DIV 11. Cell death assay was performed 24 h after OGR. Results are expressed as a percent of control (n = 5). **B.** Representative western blot overexpression of SULTA1 in mouse brain following SULT4A1 AAV injection. SULT4A1 expression levels were analyzed 21 days after injection. **C.** Quantification of fold change of exogenous SULT4A1 compared to basal levels. Data are mean ± SEM (n = 5). **D.** Representative TTC staining of AAV-GFP or AAV-SULT4A1 injected mouse brain after 24 hr of MCAO. **E.** Quantification of infarct volume obtained from **(D).** Data are mean ± SEM (n = 5). **P < 0.001 vs AAV-GFP, calculated with two-way ANOVA followed by Tukey’s post hoc test. **F.** Representative axial images from rostral to caudal of T2 MRI of stroke mice injected with GFP-AAV or SULT4A1-AAV at Day 3, Day 15, and Day 30 of reperfusion. Quantification of the brain lesion volume acquired via T2 MRI at Day 3, Day 15, and Day 30 after stroke was also shown for respective images. The infarct volume of each axial level was determined using an algorithm generated in MATLAB. Data are mean ± SEM. n = 5 for each group. *P < 0.05 and **P < 0.01, and ***P < 0.001 vs AAV-GFP. Data were analyzed using Student’s t-test.

### 7. SULT4a1 improves functional deficit after stroke in mice

Post-stroke functional decline encompasses both motor and non-motor deficits. In this study, we utilized the CatWalk system to assess motor gait abnormalities and the Elevated Plus Maze (EPM) to evaluate non-motor deficits, such as anxiety, at days 3, 8, 15, and 30 following reperfusion in MCAO mice. Anxiety and depression are among the most common non-motor complications post-stroke. The EPM is a well-established method for assessing anxiety in mice following a stroke. Our data indicate that MCAO mice with GFP control expression spent significantly more time in the closed arms of the EPM, suggesting heightened anxiety. In contrast, MCAO mice with SULT4a1 overexpression spent significantly less time in the closed arms, indicating reduced anxiety and improved non-motor function (Fig 7 A & B). Gait analysis performed using the CatWalk system showed results consistent with the EPM findings. However, on day 3 post-stroke, the MCAO mice were unable to complete a successful run on the CatWalk platform. So, the gait assessments were conducted on days 8, 15, and 30 post-stroke, measuring parameters such as average speed (Fig 7C), swing speed at day 8 (Fig 7D) and day 15 (Fig 7E), stride length at day 8 (Fig 7F), phase dispersion day 8 and 15 (Fig 7G), max contact area at day 8 (Fig 7H), and print area at day 8 (Fig 7I). Our data demonstrate that these gait parameters were significantly impaired in GFP control MCAO mice. In contrast, MCAO mice with SULT4a1 overexpression exhibited significant improvements in these gait parameters post-stroke (Fig 7). Collectively, these findings suggest that SULT4a1 overexpression provides neuroprotection and ameliorates both motor and non-motor deficits in the MCAO stroke model. **Discussion**

**Figure 7.**
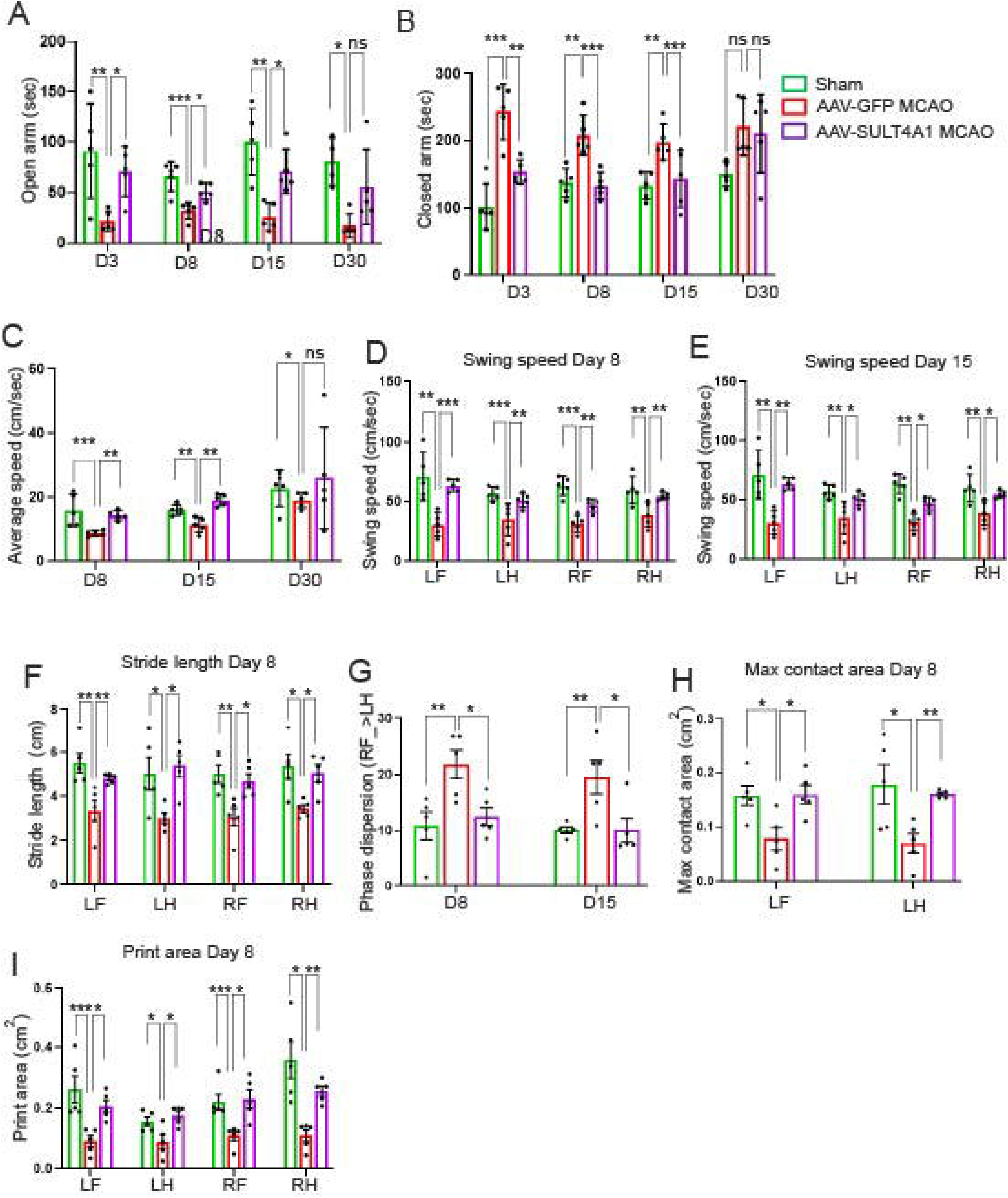
Neurological and Functional assessment of AAV-GFP and AAV-SULT4A1 injected mice after MCAO. Assessment of anxiety-like behavior via elevated plus maze analysis in sham and stroke mice stereotaxically injected with AAV-GFP or AAV-SULT4A1 at different reperfusion time points (D8 = Day 8, D15 = Day 1,5, and D30 = Day 30). Duration (sec) at **(A)** Open and **(B)** Closed arm. Data are mean ± SEM, n=5 mice for each group, Statistical analyses were performed using One-way ANOVA followed by Tuckey’s post hoc Test *p<0.05, **p <0.01, ***p<0.001. Comparison of gait analysis in sham and stroke mice stereotaxically injected with AAV-GFP or AAV-SULT4A1 at different reperfusion time points (D8 = Day 8, D15 = Day 1,5, and D30 = Day 30). Average speed (cm/sec) **(C),** Swing speed (cm/sec) of LF, LH, RF, and RH **(D)** at D8, **(E)** at D15, Stride length (cm) of LF, LH, RF, and RH at D8, **(F)**, Phase dispersion RF→LH **(G)** at D8 and D15, Max contact area (cm^2^) of LF and LH at D8 and D15, and Print area (cm^2^) **(I)** of LF, LH, RF and RH at D8,. Data are mean ± SEM, n= 5 mice for each group, Statistical analyses were performed using One-way ANOVA Bonferroni Test *p<0.05, **p <0.01, ***p<0.001. RF = right front, LF = left front, RH = right hind, and LH = left hind.

We report that SULT4a1 is neuroprotective against neuronal injury caused by OGD and ischemic stroke. The study highlights that SULT4a1 is predominantly expressed in neurons within the brain, with minimal expression in other cell types such as astrocytes. This neuronal specificity suggests that SULT4a1 likely plays an important role in regulating key functions within mature, differentiated neurons. The findings of the study point to the protective properties of SULT4a1 against neuronal damage, as SULT$a1 maintains mitochondrial function and reducing oxidative stress, both of which are critical in the pathophysiology of stroke and OGD.^7,16^

The findings of this study indicate that when SULT4a1 levels decrease in OGD or in tMCAO stroke *in vivo*, it contributes to neuronal damage, further emphasizing the importance of SULT4a1 in maintaining neuronal health. By restoring SULT4a1, the study suggests a potential therapeutic strategy for mitigating damage in stroke. Our previous study demonstrated that SULT4a1 localizes to mitochondria in neuronal cell lines like SH-SY5Y cells^7^, pointing to its role in regulating mitochondrial function. This mitochondrial localization is significant because mitochondrial dysfunction is one of the earliest events observed in both ischemic stroke and OGD-induced neuronal injury, leading to bioenergetic failure, and cell death.^24,25^ Loss of SULT4a1 levels is sufficient to cause mitochondrial defects in mouse cortical neurons indicating that SULT4a1 is playing vital role in regulating mitochondrial function via mechanisms not yet clearly understood. Mitochondrial dysfunction is pivotal to the pathophysiology of stroke^3,26^. Mitochondrial dysfunction in neurons with shRNA-mediated loss of SULT4a1 imply that loss of SULT4a1 is a critical pathophysiological process in stroke and restoring SULT4a1 levels may be a vital neuroprotective strategy. One of the key aspects of the study is the observation that SULT4a1 overexpression helps to preserve mitochondrial function and mitochondrial Δψm in neurons exposed to OGD, offering protection against mitochondrial dysfunction. Furthermore, this protective effect appears to be linked to a reduction in oxidative stress. Our findings that following OGD, neurons with SULT4a1 overexpression have lower ROS generation and decreased protein carbonylation compared to neurons with control lentiviral overexpression suggesting that SULT4a1 plays a role in mitigating oxidative damage. Given that oxidative stress is a major contributor to neuronal injury during stroke^27,28^ and other neurodegenerative diseases,^29^ this finding supports the idea that SULT4a1 could be crucial for neuronal survival under pathological conditions. However, whether SULT4a1 directly mitigates oxidative stress or indirectly by maintaining mitochondrial integrity and function remains unknown. It is likely that preservation of mitochondrial function by SULT4a1 is reducing the oxidative damage as mitochondrial dysfunction results in dysregulation of oxidative homeostasis^7^. The precise mechanisms through which SULT4a1 improves mitochondrial function and reduces oxidative stress remain unclear. The study posits that SULT4a1 may exert its protective effects through its presence in both the mitochondria and the cytosol. It could potentially be influencing mitochondrial function directly, or it may act to regulate the generation and accumulation of ROS both in the mitochondria and the cytosol. Further research will be needed to confirm whether SULT4a1 has a direct ROS-detoxifying role or whether it contributes to mitochondrial bioenergetic regulation via other pathways. Understanding the molecular interactions of SULT4a1 in these processes could offer valuable insights into its potential as a therapeutic target for stroke and other neurological conditions characterized by mitochondrial dysfunction and oxidative stress.

Overall, while this study establishes a compelling link between SULT4a1 expression and neuronal protection in stroke and OGD, additional research is necessary to fully elucidate the mechanisms by which SULT4a1 exerts its effects. It would also be beneficial to explore whether modulating SULT4a1 expression could lead to meaningful therapeutic outcomes in other animal models and, potentially, in clinical settings with future drugs that may be used to increase the expression or inhibit the loss of SULT4a1 in stroke.

## MATERIAL AND METHODS

### Primary mouse cortical neural cell culture

All experiments involving the use of animals were approved by the Institutional Animal Care and Use Committee (IACUC) at the University of Alabama at Birmingham. Primary mouse cortical neurons were prepared following the methods described previously^30,31^. In brief, cortical neurons were isolated from embryonic day 15 (E15) mouse embryos in a dissection medium containing DMEM (D6046, Millipore Sigma) and 20% horse serum (Thermo Fisher Scientific, Cat # 26050088,). The isolated cortices were digested in TrypLE (Thermo Fisher Scientific, Cat # 12605010) for 5 minutes at 37°C and triturated using a fine pipette to prepare a homogeneous single-cell suspension. The cell suspension was passed through a 40-micron cell strainer (Fisher Scientific, Cat 22-363-547) to remove cell debris. After counting, a density of 5 × 10^5 cells/mL was placed on cell culture plates pre-coated with poly-L-Ornithine (Millipore Sigma, Cat # P3655) in complete Neurobasal-A medium (Thermo Fisher Scientific, Cat # A2477501), supplemented with 10 mM glucose, 1 mM GlutaMax (Thermo Fisher Scientific, Cat # 35050061), 1 mM sodium pyruvate (Thermo Fisher Scientific, Cat # 11360070,), and B-27 (Thermo Fisher Scientific). A 50 μM 5-fluoro-2-deoxyuridine (Thermo Fisher Scientific, Cat # L16497MC) was added to the culture at DIV2 (Days in vitro) to inhibit glial cell growth. Experiments were performed with DIV11 neurons.

### Astrocytes culture

Primary astrocyte cultures were prepared from the cortices of E15 mouse embryos. The cortices were isolated and processed as described in the primary neuronal culture section, except that 5-fluoro-2-deoxyuridine was not added to the culture. The culture was grown in DMEM (Millipore Sigma, Cat # D6046) supplemented with 10% FBS (Thermo Fisher Scientific, Cat # 26050088,) for two weeks in an incubator at 37°C with 5% CO₂, with the medium replaced every three days until the culture reached confluence. The culture was then trypsinized, subcultured into a new T75 flask, and maintained until confluence. Once confluent, the culture in the flask was shaken at 240 rpm for 6 hours to remove non-astrocytic glial cells. Afterward, the astrocyte culture was washed with PBS, trypsinized, and subcultured into a new flask or on cell culture plates/cover glasses. The purity of the astrocyte culture was assessed by immunocytochemistry using glial fibrillary acidic protein (GFAP), an astrocytic marker.

### Oxygen-glucose deprivation (OGD) in primary cortical neurons

OGD in primary neurons was performed as described previously.^30,31^ In the first step, two-thirds of the Neurobasal medium from each well was collected into a Falcon tube and placed in an incubator at 37°C. The neurons were then washed twice with warm PBS to remove any residual medium. Afterward, OGD buffer [NaCl (116 mM), KCl (5.4 mM), MgSO4 (0.8 mM), NaHCO3 (26.2 mM), NaH2PO4 (1 mM), CaCl2 (1.8 mM), glycine (0.01 mM) (pH 7.4)], pre-bubbled with OGD gas (5% CO2, 10% H2, and 85% N2), was added to the neuronal culture. The culture plates were then placed in a hypoxia chamber at 37°C for 60 minutes (Biospherix Ltd., Parish, NY, USA). The hypoxia chamber was connected to the OGD gas supply to maintain an O2 level of less than 0.5% through a continuous flow of the OGD gas. An oxygen sensor (Biospherix Ltd., Parish, NY, USA) was used to monitor oxygen levels in the chamber throughout the OGD. OGD was terminated after 60 minutes to start oxygen and glucose resupply (OGR) by adding the neurobasal media collected in step first back to the cultures and placing them back into the regular incubator with normoxic conditions (5% CO2 and 95% air).

### Western blots

For western blotting, neurons were first washed with ice-cold PBS and lysed in RIPA buffer (pH 7.4, 1% Triton X-100, 0.1% SDS, 1% deoxycholate, with protease and phosphatase inhibitors). The protein concentration of each sample was determined using the BCA method. Equal volumes of lysate containing equal amounts of protein from each sample were separated by SDS/PAGE. Afterward, proteins were transferred from the gels onto a nitrocellulose membrane (Bio-Rad, Hercules, CA, USA) using the wet transfer method for 1 hour. The membranes were collected and blocked with 5% non-fat milk diluted in TBST for 1 hour at room temperature. After blocking, the membranes were incubated with the respective primary antibodies overnight at 4°C. Following overnight incubation, the primary antibodies were removed, and the membranes were washed three times with TBST (Tris-buffered saline + 0.1% Tween 20). The membranes were then incubated with HRP-conjugated anti-rabbit or anti-mouse secondary antibodies (Cell Signaling) for 1 hour at room temperature. Afterward, the membranes were washed three times (15 minutes each) with TBST. Protein signals on the membranes were detected using SuperSignal West Pico Plus Chemiluminescent Substrate (Thermo Fisher, Cat # 34578). ChemiDoc MP imaging system (Bio-Rad, USA) was used to image the membranes. To visualize β-actin, the membranes were stripped and processed for β-actin detection using an HRP-conjugated β-actin antibody (Abcam).

### Immunocytochemistry

For immunocytochemistry, neurons grown onto poly-L-Ornithine-coated glass coverslips (Carolina Biological Supply) were washed with ice-cold PBS and fixed with 4% paraformaldehyde for 15 min at room temperature. Then the neurons were washed three times with PBS and permeabilized using 0.2% Triton X-100 in PBS containing 10% donkey serum (Mllipore Sigma, Cat # D9663) for 20 min and blocked in blocking buffer (10% donkey serum in PBS) for 1 h at room temperature. Afterward, neurons were incubated in primary antibodies diluted in blocking buffer overnight at 4°C. After incubation, neurons were washed three times with PBS, and then incubated with the Alexa Fluor 488 or Alexa Fluor 555 conjugated secondary antibody (Thermo Fisher Scientific, 1:1000) diluted in PBS containing 1% (v/v) donkey serum for 1 h in the dark at room temperature. Afterward, neurons were washed with PBS three times and counterstained with DAPI (300nM) for nuclear staining. Finally, the coverslips were mounted on glass slides using mounting media (P36930, ProLong Gold, Thermo Fisher) for imaging using an LSM 710 Confocal Microscope (Carl Zeiss, Germany).

### Measurement of oxygen consumption ratio (OCR)

Oxygen consumption rate (OCR) was measured using an XFe96 Extracellular Flux Analyzer (Agilent Technologies, Santa Clara, CA, USA) following previously described protocols.^7,30,32^ Neurons were plated in a 96-well XF96 cell culture microplate (Agilent Technologies) at a density of 70,000 cells per well. OCR analysis was conducted on day in vitro (DIV) 11. Prior to the experiment, neurons were washed with 200 µL of XF DMEM medium (pH 7.4) containing 10 mM glucose, 1 mM L-glutamine, and 1 mM sodium pyruvate. The neurons were then incubated in 180 µL of the XF DMEM medium for 1 hour at 37°C. Meanwhile, the XF Sensor Cartridge, containing oligomycin, carbonyl cyanide 3-chlorophenylhydrazone (CCCP), and antimycin A + rotenone (Sigma-Aldrich), was calibrated in a CO2-free incubator. The reagents were sequentially injected into each well via the sensor cartridge to assess basal respiration, maximal respiration, and ATP turnover. The assay protocol involved a 1.5-minute mixing phase, a 1-minute wait phase, and a 2-minute measurement phase, repeated for three cycles. After completing the assay, the culture plates were collected, and protein concentration in each well was determined. Data were normalized to the protein concentration in each well.

### Mitochondrial membrane potential

Mitochondrial membrane potential in live neuronal cells was measured using the cell-permeant dye Tetramethyl rhodamine ethyl ester (TMRE; Thermo Fisher Scientific), following a previously described protocol.^7,30^ Initially, neuronal cells grown on glass coverslips were incubated with 10 nM TMRE, diluted in neurobasal medium, for 30 minutes in a CO2 incubator at 37°C. After incubation, the cells were washed twice with warm PBS, and the coverslip was placed onto a coverslip holder. A 500 µL volume of neurobasal medium was added to cover the neurons during imaging. Live-cell images were captured using an LSM710 confocal microscope (Zeiss, Germany) with a temperature-controlled stage-top incubator, at 30-second intervals. After recording a baseline, 20 µM CCCP (Sigma-Aldrich) was added, and images were captured for an additional 2 minutes. Zen software (Zeiss, Germany) was used to quantify TMRE intensity as a relative ratio before and after CCCP addition (ΔF/F₀).

### Measurement of oxidative stress

Oxidative stress in neurons was assessed using the CellROX Green Reagent (Thermo Fisher Scientific), following the manufacturer’s instructions. Neurons grown on glass coverslips were incubated with 5 µM CellROX Green Reagent for 30 minutes at 37°C in a CO2 incubator. After incubation, the cells were washed with PBS. Live-cell images were captured using an LSM710 confocal microscope (Zeiss, Germany). Zen software (Zeiss, Germany) was used to quantify oxidative stress as relative fluorescence density.

### Stereotaxic injection

SULT4A1 Adeno-associated virus (AAV) or Control AAV were delivered into the cortex of anesthetized mice (isoflurane anesthesia induction chamber, 3% isoflurane, 97% air, 250 CC/min) by stereotaxic injection.^31^ Two injections of AAV9 particles (1 μl of ∼10^13^ titer) were made in the cortex using coordinates, 1.0 mm rostral, 1.5 mm lateral, and 1.5 −3.0 mm ventral from bregma, and 1.0 mm caudal, 1.5 mm lateral, and 1.5-3.0 mm ventral from bregma with a 26 G Hamilton syringe mounted on a stereotaxic stand (Kopf). Ventrally, we injected first (0.5µl at 3.0mm, after a 3 min wait, retracted the needed to 1.5 mm location and injected the rest 0.5µl. The virus particles were delivered using an automatic pump (Stoelting Co., IL, USA) at a speed of 0.5 μl/min. The syringe remained inserted for 5 min after completion of injection and was retracted slowly (0.5 mm/min) to avoid backflow of injected virus. The MCAO surgery on the mice was performed three weeks after AAV virus injection.

### Middle Cerebral Artery Occlusion (MCAO)

Two-month-old CD1 mice underwent MCAO surgery following a previously described method.^31^ The surgery was performed under 3% isoflurane in medical-grade oxygen. The mice’s body temperature was monitored using a rectal thermistor probe and maintained at 37°C throughout the procedure with the aid of a thermostatically regulated heating pad (CMA/150, CMA/Microdialysis). To occlude the right middle cerebral artery (MCA), a silicone-coated 7-0 nylon monofilament (Doccol Corporation, MA, USA) was inserted into the right internal carotid artery (ICA) through a small incision in the right external carotid artery. The monofilament was then advanced through the ICA to the point where the anterior carotid artery begins, blocking blood flow to the MCA. Successful occlusion was confirmed using a Laser-Doppler flowmetry probe to monitor the blood flow blockage in the MCA territory. Mice with less than 80% reduction in cerebral blood flow in the MCA territory were excluded from the study. After 60 minutes of occlusion, the filament was slowly retracted from the ICA to initiate reperfusion. The sham-operated mice underwent the same surgical procedure, but without the insertion of the monofilament. The MCAO mice were used at various time points for biochemical analysis, immune-staining, or for longitudinal MRI and functional analysis.

### TTC Staining of mouse brain

The infarct volume was calculated 24 hours after reperfusion following tMCAO using 2,3,5-Triphenyltetrazolium chloride (TTC) staining. The mouse brain was dissected and rinsed in ice-cold PBS to remove any residual blood. It was then placed at −20°C for 10 minutes before being cut into 2mm thick coronal sections using razor blades and a mouse brain matrix. This brief step in −20°C helps in smooth cutting of the sections. The sections were incubated in a 1% TTC solution, diluted in PBS, for 20 minutes at 37°C. After incubation, the sections were fixed in 4% paraformaldehyde and imaged using a camera or scanner. ImageJ software was used to calculate the infarct volume according to the previously described formula (I_EA_ = (L − N)/L*100; where I = absolute unadjusted infarct area; I_EA_ = edema-adjusted I; L = left (contralateral) hemisphere area; N = non-infarcted tissue in the ipsilateral region).^33^

### Magnetic Resonance Imaging

MRI is a non-invasive and translationally relevant tool for brain lesion assessment.^34,35^ T2/FLAIR imaging was conducted on MCAO mice under 1%-3% isoflurane anesthesia, with temperature control maintained at 37.5°C, using a 9.4T Bruker Biospin scanner. Data collection will occur at 3, 7, and 15 days post-MCAO or sham surgery. Infarct volumes are quantified using semi-automated segmentation with ITK-SNAP software (version 4.0.2) on T2/FLAIR-weighted images acquired at 9.4T Bruker Biospin scanner. Hyperintense regions corresponding to ischemic tissue were identified based on signal intensity thresholds and then manually refined by a trained observer. The final infarct volume was reported in mm³.

### Elevated Plus-Maze

The Elevated Plus Maze (EPM) test is commonly used to assess anxiety-like behavior. The maze consists of two intersecting arms, one with an open platform and the other with a closed platform. The arms are elevated 20 cm above the ground, with each arm measuring 6 cm in width and 75 cm in length (Noldus, Wageningen, The Netherlands). For this test, mice were placed at the center of the maze and allowed to explore for 5 minutes. Their voluntary movements, including the time spent in each arm and the open central area, were recorded. The time spent in the open and closed arms was analyzed to assess anxiety levels using Ethovision software (Ethovision 11, Noldus Information Technologies, Leesburg, VA, USA)

### Gait analysis

Gait analysis was performed following the manufacturer’s instructions and a previously described protocol using an automated computer-assisted system (CatWalk™ XT10, Noldus Information Technology, Wageningen, The Netherlands).^36^ The system was placed in a dark (<20 lux), quiet room. Experimental animals were trained to walk from one end to the other of a 1.3-meter-long glass platform two weeks prior to the final experiment. After completing the training, the mice underwent sham or tMCAO surgery. Gait analysis was performed on these animals at 8-, 15-, and 30-days post-surgery. On the day of the experiment, the mice were placed at one end of the glass platform and encouraged to walk across. Each mouse completed three practice walks before the final measurements were taken. A high-speed camera positioned underneath the glass platform recorded each full walk. At least three valid runs were required, each lasting between 0.5 and 5 seconds, with a maximum speed variation of 60%. Disrupted walks, where the mouse stopped or walked backward, were excluded from the analysis. After adjusting the intensity threshold, footprints were identified and labeled to enable the analysis. A broad range of gait parameters, including run duration, average speed, swing speed, stride length, phase dispersion, maximum contact area, print area, and others, were assessed.

### Statistical analysis

Neuronal cultures were randomly distributed into control and experimental groups. The number of the mice used in the study was determined by power analysis to obtain statistical significance. Western blot and fluorescence images were quantified using ChemiDoc (Bio-Rad) and Zen software, respectively. Statistical analyses were performed using GraphPad Prism version 9 (GraphPad, USA). Quantified data are presented as mean ± SEM. One-way analysis of variance (ANOVA) with Tukey’s post hoc test was used for multiple group comparisons or Student’s t-test for two group comparison. P<0.05 were considered statistically significant for the analyses.

## Acknowledgments

This work was supported by grants from the NIH, R01NS119479 (SAA), R21AG084284 (SAA)

